# Understanding Genome Structure Facilitates the Use of Wild Lentil Germplasm for Breeding: A Case Study with Shattering Loci

**DOI:** 10.1101/2023.12.18.572160

**Authors:** Zhe Cao, Didier Socquet-Juglard, Ketema Daba, Albert Vandenberg, Kirstin E Bett

**Affiliations:** Department of Plant Sciences, University of Saskatchewan. 51 Campus Dr. Saskatoon, SK S7N 5A8 Canada

**Keywords:** pod dehiscence, interspecific hybridization, *Lens culinaris*, *Lens orientalis*, *Lens odemensis*, crop wild relatives

## Abstract

Plant breeders are generally reluctant to cross elite crop cultivars with their wild relatives to introgress novel desirable traits due to associated negative traits such as pod shattering. This results in a genetic bottleneck that could be reduced through better understanding of the genomic locations of the gene(s) controlling this trait. We integrated information on parental genomes, pod shattering data from multiple environments, and high-density genetic linkage maps to identify pod shattering QTLs in three lentil interspecific recombinant inbred line (RIL) populations. The broad-sense heritability on a multi-environment basis varied from 0.46 (in LR-70, *Lens culinaris x L. odemensis*) to 0.77 (in LR-68, *L. orientalis x L. culinaris*). Genetic linkage maps of the interspecific populations revealed reciprocal translocations of chromosomal segments that differed among the populations, and which were associated with reduced recombination. Segregation distortion was also observed for clusters of SNPs on multiple chromosomes per population, further affecting introgression. Two major QTL, on chromosomes 4 and 7, were repeatedly detected in the three populations and contain several candidate genes. These findings will be of significant value for lentil breeders to strategically access novel superior alleles while minimizing the genetic impact of pod shattering from wild parents.

## INTRODUCTION

Pod shattering, or dehiscence, is a mechanism whereby mature seed pods split to disperse seeds into the environment. This characteristic is crucial for wild species to efficiently disperse their seeds but is clearly undesirable in crop production. In legumes, the fruit is a pod with seeds along a ventral suture (Esau 1977). At maturity, the pods dehisce, open explosively, and expel seeds out into the surrounding environment. Selection for non-shattering phenotypes began at the very early stage of crop domestication (Meyer and Purugganan 2013). Genetic analysis of genes that control shattering has been conducted in various legumes including soybean, lupin, pea, cowpea, and common bean (reviewed by Ogutcen et al. 2018). Various results indicated that the trait is controlled by a single gene, a set of two genes, or multiple genes depending on the population being examined. In lentil, shattering was originally characterized as a monogenic trait (Tahir and Muehlbauer 1994), but later it was suggested to be polygenic based on continuous distribution of data in progeny of a *L. culinaris* x *L. orientalis* population (Fratini et al. 2007).

In lentil breeding, the development of new cultivars with improved yield and quality has been a primary goal to address both the requirements of farmers and the expectations of the market. Due to intensive selection for increased crop productivity and uniformity for decades, valuable genes controlling other traits that may be valuable for future breeding, or for the adaptation of the crop to changing environments, have become lost in elite cultivars and breeding lines. The limited genetic variation present in superior breeding germplasm may hinder the development of cultivars capable of meeting future needs. Wild species offer a source of alleles absent in the cultivated gene pool. In lentil, studies have shown that wild lentil accessions possess a range of novel beneficial alleles related to cold hardiness, drought tolerance, and disease resistance (Ali et al. 1999; Gupta et al. 2006; Bhadauria et al. 2017). Plant breeders, however, are often reluctant to use wild relatives considering that they may harbor numerous deleterious genes that outweigh the beneficial effects of desirable genes. The introgression of a desired gene from a wild accession could result in introgression of a sizeable segment of wild chromosome containing many potentially undesirable genes that even decades of traditional breeding could not be effectively recombined (Tanksley and Nelson 1996). Understanding the genomic relationships between a crop and its wild relatives allows for strategic hybridization and selection to incorporate positive alleles while avoiding negative ones.

One of the major concerns of using wild lentil relatives or species would be simultaneous introduction of gene(s) that cause pod dehiscence. In lentil, Tahir and Muehlbauer (1994) used pod indehiscence (*Pi*) and other morphological markers in conjunction with restriction fragment length polymorphisms (RFLP) and alloenzymes markers to construct a linkage map of *L. culinaris* × *L. orientalis*. They found that genes controlling *Pi* and β-galactosidase were genetically linked. By using simple sequence repeat (SSR) markers and a recombinant inbred line (RIL) population from *L. culinaris* ‘Lupa’ x *L. orientalis* ‘BGE 016880’, Fratini et al. (2007) identified three shattering QTLs on three independent linkage groups (their LGII, LGIII, and LGV). The shattering QTL on their LGIII was in proximity to QTL associated with early growth rate (number of branches at the 1^st^ node and height of 1^st^ node). The unknown genomic position of these markers makes it difficult to compare them or to link them to other populations or to sequenced *Lens* spp. genomes. Recently, Chen (2018) used a SNP map of a *L. culinaris* cv. ‘Eston’ x *L. ervoides* acc. IG 72815 (LR-26) RIL population to identify a single locus associated with dehiscence that mapped to the region around 430 Mb on chromosome 7 of the Lcu2.RBY lentil genome (Ramsay et al. 2021). A reduced level of recombination was also observed on that chromosomal segment, resulting in genetic linkage between the dehiscence QTL and another QTL that controls anthracnose resistance. Those results suggest the using wild species to introgress anthracnose resistance could bring along a pod shattering gene as well, which would be problematic in genetic enhancement of elite cultivars.

The objective of this study was to advance our knowledge of the genetic control of pod shattering in lentil species by examining whether the responsible dehiscence genes consistently map to the same chromosomal segments across diverse genetic backgrounds, by assessing the extent of chromosomal segments linked with pod shattering QTLs, and by determining how this differs across several different interspecific cross combinations. We used three RIL populations derived from crosses between *L. culinaris* and *L. orientalis* (LR-68 and LR-86) or *L. odemensis* (LR-70), and phenotyped them for dehiscence over multiple years and locations. We used genetic linkage maps associated with sequenced genomes to detect and localize QTLs controlling pod shattering. The information gained from this study will provide breeders with tools and a strategy for crossing lentil with wild relatives to strategically limit the negative effect of pod dehiscence while introgressing useful genetic variation.

## MATERIAL AND METHODS

### Plant material

Three F_8_-derived interspecific RIL populations (LR-68, LR-70, and LR-86), developed using single seed descent, were used in this study. LR-68 (n=120 RILs) was derived from a cross between IG 72643 (*L. orientalis* Boiss.) and the breeding line 3339-3 (*L. culinaris* Medik.). LR-70 (n=112 RILs) was derived from a cross between ‘Eston’ (*L. culinaris*) and IG 72623 (*L. odemensis*). LR-86 (n=93 RILs) was derived from the cross between ‘Lupa’ (*L. culinaris*) and BGE 016880 (*L. orientalis*) (Fratini et al. 2007; Yuan et al. 2021). All wild parents were dehiscent and cultivated ones were indehiscent.

Field trials, set up in a randomized complete block design with three replicates, were established at four field locations: Crop Science Field Lab Seed Farm (CSFL), Investigation, Sutherland, and Preston, in 2017 or 2018, all near Saskatoon, SK, Canada. Twelve plants were arranged in one plot representing one replication for each genotype, with 30 cm between each plot. Seeds were scarified using sandpaper and sown to maintain 30 cm between plants. Standard field agronomic practices for cultivated lentil were used to maintain plots.

When the first pods were visible in the field, the three largest plants per plot were each enclosed in mesh bags to capture any seeds from dehisced pods. Bags were left open at the top of the plants but were closed at the base to collect all seeds. When 50 to 75 % pod maturity was reached, mesh bags were closed and individually harvested. Plants were dried at 32°C and stored in a dry container prior to rating for dehiscence. Seeds of each replicate that were found loose in the mesh bags were separated from the plant material using an air blower, weighed, counted, and considered as shattered seeds. The seeds that could only be separated after mechanical threshing were considered non-shattered seeds and were counted and weighed separately. A value for percent dehiscence was calculated for each line as the ratio of the total number of seeds shattered divided by the total number of seeds collected for the individual plants. The results from the three plants sampled in each plot were averaged to give a value for each plot.

### DNA extraction, library construction, sequencing, and SNP calling

Leaf tissue was collected and immediately frozen in liquid nitrogen and stored at -80°C before DNA extraction. Genomic DNA was extracted by using a modified CTAB method (Doyle and Doyle 1990). The quantity and quality of DNA was examined on 1% agarose gel and Quan-iT PicoGreen DsDNA assay kit (Life Technologies, Carlsbad, CA, USA) on a FLUOstar Omega fluorometer (BMG Labtech, Ortenberg, Germany).

For RILs and parents of the LR-68 and LR-70 populations, library construction was performed based on a two-enzyme system of Pstl and Mspl to cut genomic DNA following the protocol described by Poland and Rife (2012). DNA fragments were sequenced on an Illumina HiSeq 2500 sequencer (Illumina, San Diego, CA, USA) using the paired-ended mode (2 × 100 bp) at the DNA Sequencing Laboratory, NRC-Saskatoon. Raw sequencing reads were demultiplexed and barcode sequences removed before filtering and trimming using Trimmomatic using the default settings (Bolger et al. 2014). The trimmed reads were aligned against the *L. culinaris* reference genome Lcu.2RBY (Ramsay et al. 2021) using Bowtie allowing only end-to-end matches (Langmead and Salzberg 2012). SNP calling was performed using mpileup in Samtools (Li et al. 2009). The VCF format output was then converted to ABH format for the ease of linkage map construction. Library construction, sequencing and SNP calling of the LR-86 population was similar and is described in detail in Yuan et al. (2021).

### Phenotypic data analysis

The frequency distribution of raw pod shattering data was initially visualized for the LR-68, LR-70, and LR-86 populations across various site-years to determine if any noticeable skewness in the data distribution was present. To determine whether data skewness could be reduced, a square-root transformation was performed on the raw shattering data. Whichever dataset, either the raw or the transformed, that demonstrated the lowest level of skewness was chosen for subsequent data analyses. Pearson’s correlation was conducted to assess the consistency of pod shattering performance across various site-years for each population. Heritability (*H^2^)* was calculated using the statistical model: Y_ijk_ = µ + G_i_ + E_j_ + G_i_ × E_j_ + ε_ijk_, where Y_ijk_ represents the measured pod shattering for the plant_ijk_, µ is the population mean, G_i_ is the genetic effect, E_j_ is the environment effect (site-year), G_i_ × E_j_ is the interaction between genotype and environment, and ε_ijk_ is the random error. All components in this model were treated as random effects. All analyses were conducted using statistical software JMP 13 (SAS Institute, Cary, NC, USA).

### Linkage map construction, QTL analysis and search for candidate genes

Genetic linkage maps for all three populations were first constructed from SNP data with the exclusion of those markers with significantly distorted segregation (p < 0.01). MSTmap (Wu et al. 2008) was used to develop a linkage map with the following parameters: cut_off_p_value: 1e-6, missing_threshold: 0.40, distance function: kosambi, no_map_dist: 15.0, no_map_size: 0, estimation_before_clustering: yes, and detect_bad_data: yes. IciMapping (version 4.2.53) was then used to correct the order of SNPs within each linkage group (LG) using the setting of threshold value (LOD = 5), k-Optimality algorithm by REC (2-OptMAP) and Shortest NN (10) (Meng et al. 2015). Rippling of markers was conducted in the same software with the setting of Window Size = 5 by REC. Using the markers without segregation distortion to form the backbone of the linkage groups, we incorporated previously excluded distorted markers for linkage map reconstruction. Including distorted markers improved the consistency of linkage maps with the genome and coverage of the genome. Genetic linkage map data can be accessed here: https://knowpulse.usask.ca/study/shattering-loci-case-study.

Pairwise recombination frequency for all pairs of SNPs was visualized by heatmap in the R package R/qtl (Broman et al. 2003). The visualization of synteny between linkage maps and physical chromosomes (wild and cultivated parents) and allele frequency (origin of allele) (RILs) was conducted in the web-based platform SynVisio (https://genomevis.usask.ca/linkage-map/#/). Genome assemblies used included Lcu.2RBY (Ramsay et al. 2021), Lor.1WPS (*L. orientalis* acc. BGE 016880) (Ramsay et al. 2022a) and Lod.1TUR (*L. odemensis* acc. IG 72623) (Ramsay et al. 2022b).

Prior to QTL analysis, SNPs that had identical segregation patterns were combined into bins along each LG using a custom Python script (Python 3.8) (https://github.com/cjun01/linkage_map_binning). The recombinant breakpoints were the boundaries of adjacent bins. QTL mapping was implemented using these binned maps and pod shattering data via the MET method in iciMapping. This approach is used to investigate QTL-environment interaction where the QTL average effect and QEI effects could be properly estimated (Li et al. 2015). Inclusive composite interval mapping (ICIM) was used to identify and localize QTLs. The LOD threshold for declaring a marker as significant was determined by permutation test of 1000 iterations with the cut-off value set at 95^th^ percentile of LOD scores. The anchor of possible QTL regions on physical chromosomes was conducted by plotting boundaries of QTLs (“LeftMarker” and “RightMarker”) output from IciMapping.

To identify candidate genes falling into QTL regions, the keywords “shattering” and “indehiscence” were used to search all possible protein sequences in the NCBI database, resulting in 1730 sequences. After removal of redundant sequences using the software CD-hit version 4.81 (Fu et al. 2012), 48 protein sequences remained (Table S1). We conducted a reciprocal BLASTp search by blasting the annotated genes from the Lcu.2RBY genome assembly against the 48 reference proteins and vice versa. An e-value of 1-e^10^ was used as the threshold to consider the matches significant. The identified cultivated genes from the Lcu.2RBY genome were subsequently cross-referenced via BLASTn to find the corresponding wild alleles in the Lor.1WPS, Lod.1TUR, Ler.1DRO (*L. ervoides* acc. IG 72815; Ramsay et al. 2021), and Ler.1DRT (*L. ervoides* acc. L01-827a; unreleased internal version) genomes. The *L. ervoides* accessions, L01-827a and IG 72815, are additional, sequenced, wild lentils that exhibit pod shattering. Sequences of shattering gene homologs within shared QTL regions among LR-68, LR-70, and LR-86 were extracted from the Lcu.2RBY, Lor.1WPS, Lod.1TUR, Ler.1DRO, and Ler.1DRT assemblies and subjected to a multiple sequence alignment of their protein sequences and the 1000 bp upstream regulatory gene sequences to detect sequence variation between the cultivated and wild forms.

## RESULTS

### Quantitative segregation for shattering across all RIL populations

All three populations displayed quantitative distributions for percent shattering at all site-years (Fig. 1; Table S2). The cultivated parents always had mean percent shattering values below 10 % except for ‘Eston’ from the 2018 Preston (16.2 %) and 2018 Sutherland (15.5 %) sites. The wild parent shattering values were always in the range of 75 to 98 %, except ‘IG 72643’ from the 2018 Sutherland site (58 %). Some site-years were more skewed than others, generally towards less shattering – e.g., LR-70 in 2017 at the CSFL site and in 2018 at the Sutherland site.

**Figure 1.**
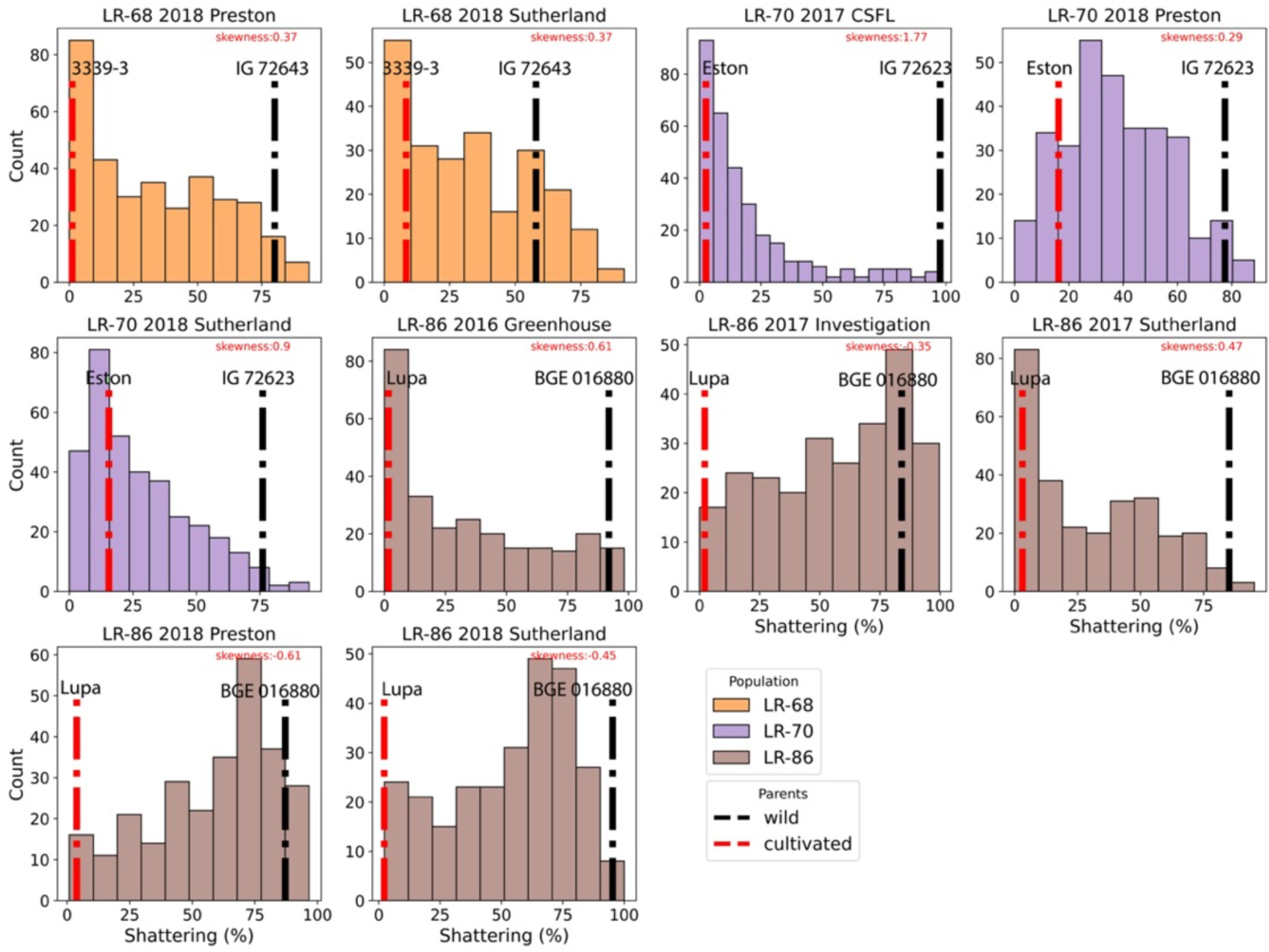
Distributions of percent pod shattering results for lentil interspecific RIL populations LR-68, LR-70, and LR-86 populations across six environments. Black vertical lines indicate values for the wild parent and red vertical lines indicate the cultivated parent in each environment.

A square-root transformation of raw shattering data was performed to determine if skewness could be decreased (Fig. S1). The shattering data after transformation exhibited reduced skewness in the LR-68 and LR-70 populations, but increased skewness in the LR-86 population. Therefore, for the subsequent statistical analyses, transformed data were used for LR-68 and LR-70, while non-transformed data were used for LR-86.

Significant correlations were observed for pod shattering among environments for all three populations (Figure S2 A, B, C). Based on the variance component partition analysis, no environment or genotype × environment effect was observed for LR-68 (Figure S2D and Table S3). However, in LR-70 and LR-86, the environment component explained 20 - 24 % of total variation, and there was a genotype × environment interaction that accounted for approximately 10 % of total variation. Broad-sense heritability (*H*^2^) ranged from high in LR-68 (0.77) to moderate in LR-70 (0.46) and LR-86 (0.54).

### Linkage maps

The 2442 SNPs for LR-68 (*L. orientalis x L. culinaris*) were anchored into six LGs spanning 1233 cM. LG1, LG3, and LG4 were named based on the corresponding chromosomes of the *L. culinaris* reference genome, Lcu.2RBY (Fig. 2). SNPs associated with both chromosomes 2 and 5 grouped together in LG2 due to pseudolinkage (Fig. 2; Fig. S3A). Meanwhile, LG5 and LG6 corresponded to chromosomes 6 and 7 of the *L. culinaris* reference genome Lcu.2RBY, respectively. Segregation distortion was observed for most SNPs on LG4 of this map, with the alleles predominantly originating from the wild parent, IG 72643 (Fig. 2). A review of the recombination frequencies and LOD scores among all SNP pairs confirmed areas with decreased recombination and instances of pseudo-linkage, such as that seen in LG2 (Fig. S3A). Reduced recombination was also observed in several large chromosomal segments of other linkage groups including LG1, LG5 and LG6. Binning of SNPs based on co-segregation resulted in 775 bins of which 357 each containing a single unique marker and the rest multiple markers. Binned and un-binned segregation matrices are available here: https://knowpulse.usask.ca/Geneticmap/3680022.

**Figure 2.**
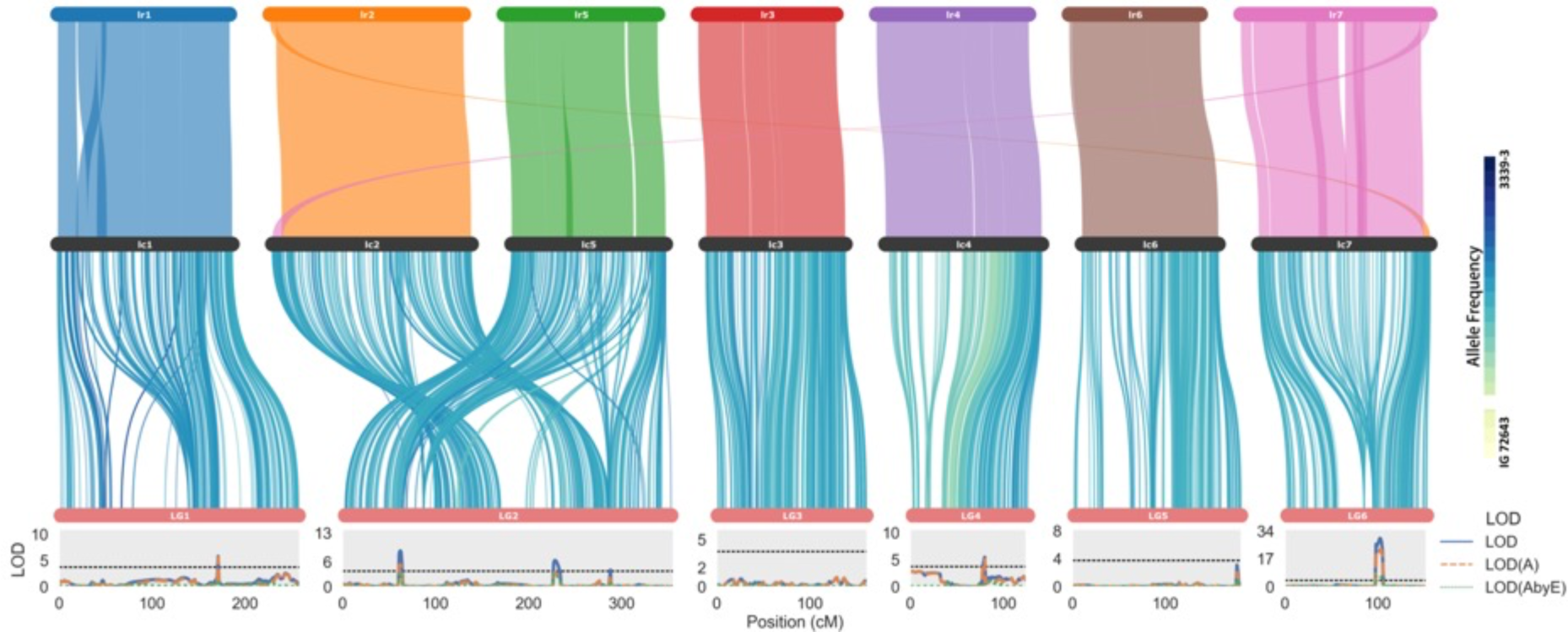
Overview of the associations among *L. orientalis* (Lor.1WPS; acc. BGE 016880; lr) and *Lens culinaris* (Lcu.2RBY; acc. CDC Redberry; lc) reference genomes, genetic linkage map of RIL population LR-68 (*L. orientalis* acc. IG 72643 *x L. culinaris* breeding line ‘3339-3’), and shattering QTLs in this population. Top ribbon shows the synteny blocks between the reference genomes. Mid panel ribbon displays the correspondence between the *Lens culinaris* genome and the LR-68 genetic linkage map. Ribbon lines between the genome and genetic linkage map are shaded according to the level of allele frequency bias towards the cultivated parent “3339-3” (dark blue) or the wild parent “IG 72643” (light green). Bottom panel shows QTL LOD scores of pod shattering along the LR-68 genetic linkage map, with dotted lines representing the LOD threshold.

For LR-70 (*L. culinaris x L. odemensis*), the 3927 SNPs that passed quality control were mapped into four LGs spanning a total genetic distance of 1069 cM. SNPs in Lcu.2RBY chromosomes 1 and 5 were grouped under LG2, as distinguishing them into separate LGs was not possible (Fig. 3). In addition, LG2 consisted of SNPs that were also associated with Lcu.2RBY chromosomes 2, 6, and 7. LG3 and LG4 were numbered corresponding to Lcu.2RBY chromosomes 3 and 4 where the SNPs were positioned. Segregation distortion was evident on LG1 and LG2 (Fig. 3). In the former, distortion mainly originated from regions corresponding to Lcu.2RBY chromosome 5 where a larger portion of the progeny inheriting alleles from the wild parent, IG 72623. In the latter, distortion was more likely to occur in regions associated with Lcu.2RBY chromosomes 2, 7 and the distal end of 6, where offspring tended to inherit alleles more frequently from the cultivated parent, cv. Eston. Extensive regions of reduced recombination were observed in LG1 and LG2 (Fig. S3B). The binning of co-segregating SNPs within each LG resulted in a total of 616 bins of which 207 were single markers. Binned and un-binned segregation matrices are available here: https://knowpulse.usask.ca/Geneticmap/3680021.

**Figure 3.**
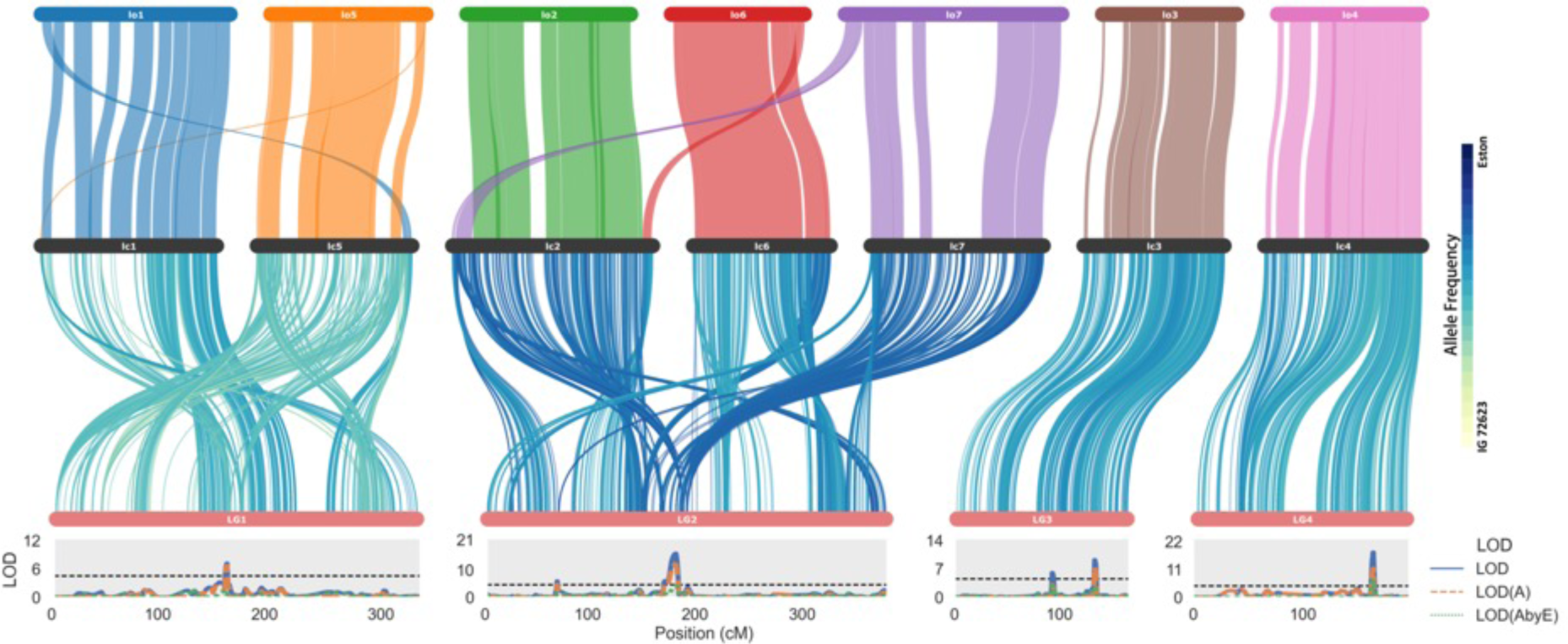
Overview of the associations among *Lens odemensis* (Lod.1TUR acc. IG 72623; lo) and *Lens culinaris* (Lcu.2RBY; acc. CDC Redberry; lc) reference genomes, genetic linkage map of RIL population LR-70 (*L. culinaris* cv. ‘Eston’ x *L. odemensis* acc. IG 72623), and shattering QTLs in this population. Top ribbon shows the synteny blocks between the reference genomes. Mid panel ribbon displays the correspondence between the *Lens culinaris* reference genome and the LR-70 genetic linkage map. Ribbon lines between the genome and the genetic linkage map are shaded according to the level of allele frequency bias towards the cultivated parent “Eston” (dark blue) or the wild parent “IG 72623” (light green). Bottom panel shows QTL LOD scores of pod shattering along the LR-70 genetic linkage map, with dotted lines representing the LOD threshold.

For LR-86 (*L. culinaris x L. orientalis*), a total of 2642 non-distorted markers were used for linkage map construction, resulting in six LGs that spanned 1051 cM. Other LGs were numbered based on where their SNPs were mapped on Lcu.2RBY chromosomes (Fig. 4). SNPs from chromosomes 2 and 7 were pseudo-linked (Fig. 4; Fig. S3C) in this map. Much like LR-68 and LR-70, the regions on LG2 experiencing chromosomal exchange also exhibited diminished recombination (Fig 4; Fig. S3C). After merging the co-segregating SNPs into SNP bins, 637 bins were created in total, out of which 245 represented single markers. Binned and un-binned segregation matrices are available here: https://knowpulse.usask.ca/Geneticmap/3680020.

**Figure 4.**
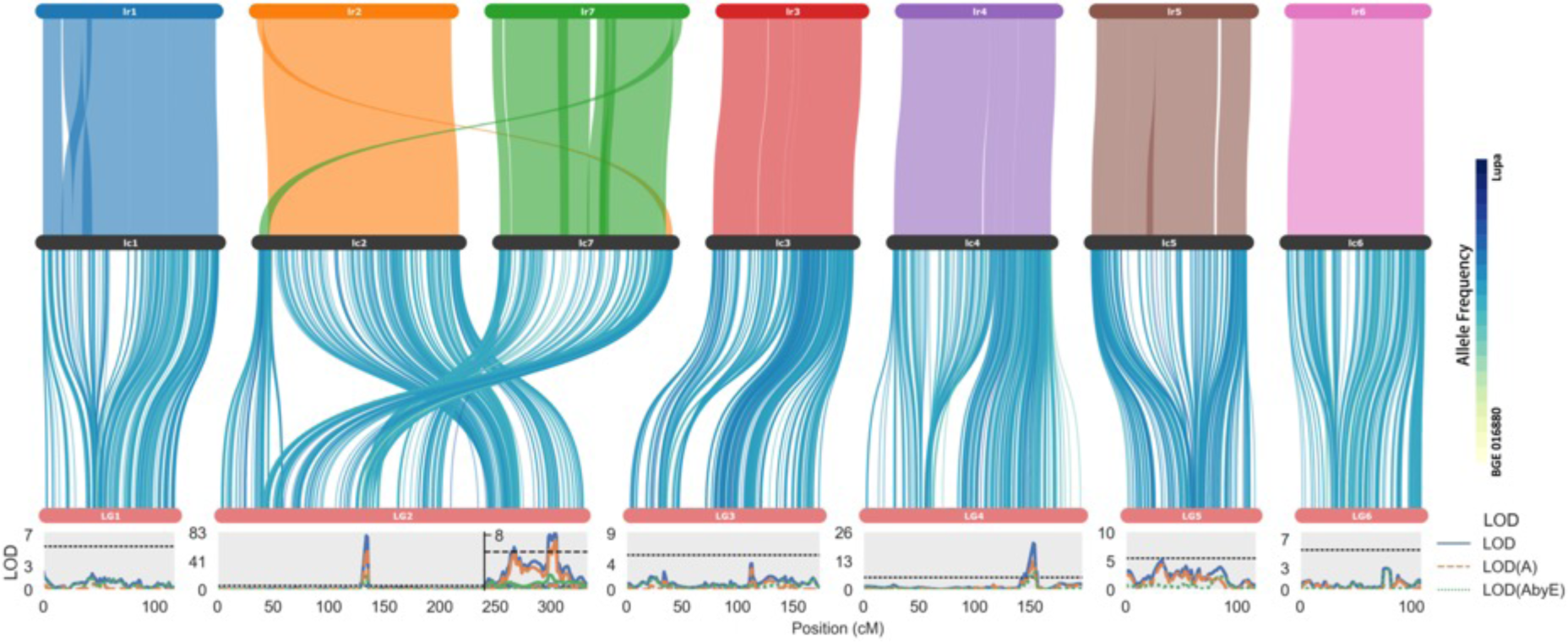
Overview of the associations among *L. orientalis* (Lor.1WPS; acc. BGE 016880; lr) and *Lens culinaris* (Lcu.2RBY; acc. CDC Redberry; lc) reference genomes, genetic linkage map of RIL population LR-86 (*Lens culinaris* cv. ‘Lupa’ x and *L. orientalis* acc. BGE 016880), and shattering QTLs in this population. Top ribbon shows the synteny blocks between the reference genomes. Mid panel ribbon displays the correspondence between the genome and LR-86 genetic linkage map. Ribbon lines between the genome and the genetic linkage map are shaded according to the level of allele frequency bias towards to the cultivated parent “Lupa” (dark blue) or the wild parent “BGE 016880” (light green). Bottom panel shows QTL LOD scores of pod shattering along the LR-86 genetic linkage map, with dotted lines representing the LOD threshold. The y-axis for the QTL plot at the right-hand end of LG2 is zoomed in to show the smaller QTL relative to the large one to the left.

### Identification of shattering QTLs

In LR-68, using the binned map and square-root normalized pod shattering data from two environments, five QTLs were identified (Fig. 2 bottom). Four of these (*qPS.LR-68_LG1.1*, *qPS.LR-68_LG2.1*, *qPS.LR-68_LG4.1*, and *qPS.LR-68_LG6.1*) remained significant after partitioning out environmental effects, maintaining LOD(A) values above the threshold (Table S4). The phenotypic variation explained (PVE) by these QTLs ranged from 2.81 % to 26.89 %. All of them had the wild allele associated with increased pod shattering.

In LR-70, the QTL analysis using the binned map and square-root normalized pod shattering data from three different environments yielded six significant QTLs (Fig. 3 bottom). Five of them (*qPS.LR-70_LG1.1*, *qPS.LR-70_LG2.2*, *qPS.LR-70_LG3.1*, *qPS.LR-70_LG3.2*, and *qPS.LR-70_LG4.3*) remained significant after removing the environment effect (Table S4). The PVE effects of these QTLs ranged from 4.18 % to 21.82 %, and the wild alleles from all of them contributed to increased pod shattering.

In LR-86, the joint analysis between the binned map and raw pod shattering data from five different environments resulted in the identification of four QTLs, among which three (*qPS.LR-86_LG2.1*, *qPS.LR-86_LG2.5*, and *qPS.LR-86_LG4.2*) were still significant after removal of the environment effect (Fig. 4 bottom; Table S4). Their PVE effects ranged from 3.30 % to 40.05 %. Alleles of these QTLs accounting for increased pod shattering were from the wild parent.

Across all three populations, QTLs consistently co-localized on the specific genomic regions of Lcu.2RBY chromosomes 4 (from 434 to 442 MB) and 7 (from 436 to 473 MB) (Fig. 5). The QTLs (*qPS.LR-68_LG6.1*, *qPS.LR-70_LG2.2*, and *qPS.LR-86_LG2.1*) on Lcu.2RBY chromosome 7 had larger effects (PVE = 14.91-40.05 %) than other QTLs (*qPS.LR-68_LG4.1*, *qPS.LR-70_LG4.1*, and *qPS.LR-86_LG4.1*) on Lcu.2RBY chromosome 4 (PVE= 2.82 – 21.86 %). Long QTL intervals were determined for those on Lcu.2RBY chromosome 7 in LR-70 and LR-86 but not LR-68. When evaluating the effects of QTLs on Lcu.2RBY chromosomes 4 and 7, offspring with a wild allele at both loci showed the most pronounced pod shattering (Fig. 6). This was followed by those with one cultivated and one wild allele, which had moderate pod shattering, and then by those with two cultivated alleles, which exhibited the least amount of pod shattering.

**Figure 5.**
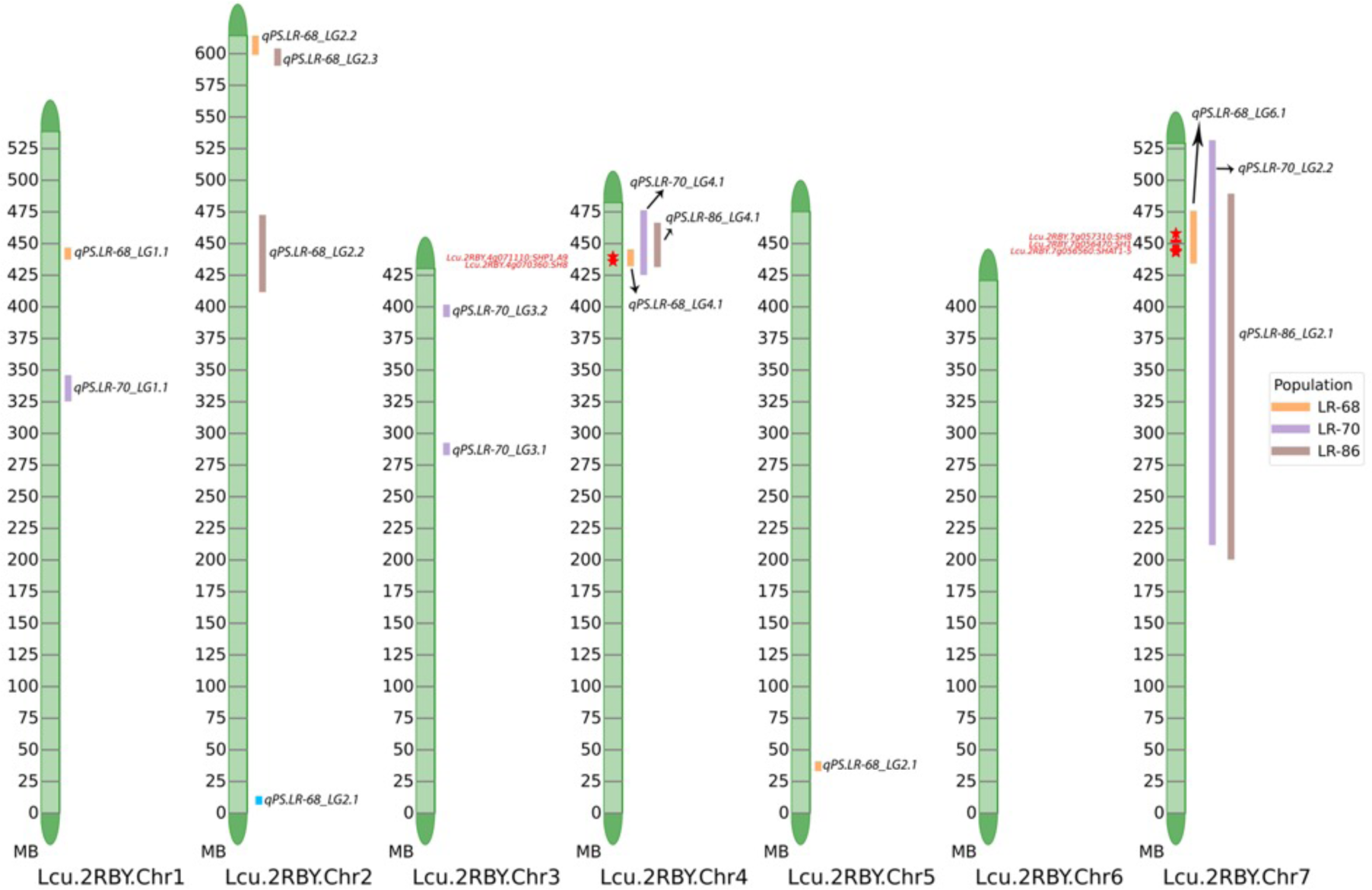
Projection of QTL intervals and candidate genes for shattering identified in the LR-68, LR-70, and LR-86 populations on the reference genome (Lcu.2RBY) of *Lens culinaris*. QTL intervals are represented by orange (LR-68), purple (LR-70), and brown columns (LR-86) to the right of each chromosome. Candidate shattering genes are plotted in the format of “Lcu.2RBY gene: known shattering gene homolog” on the left of each chromosome.

**Figure 6.**
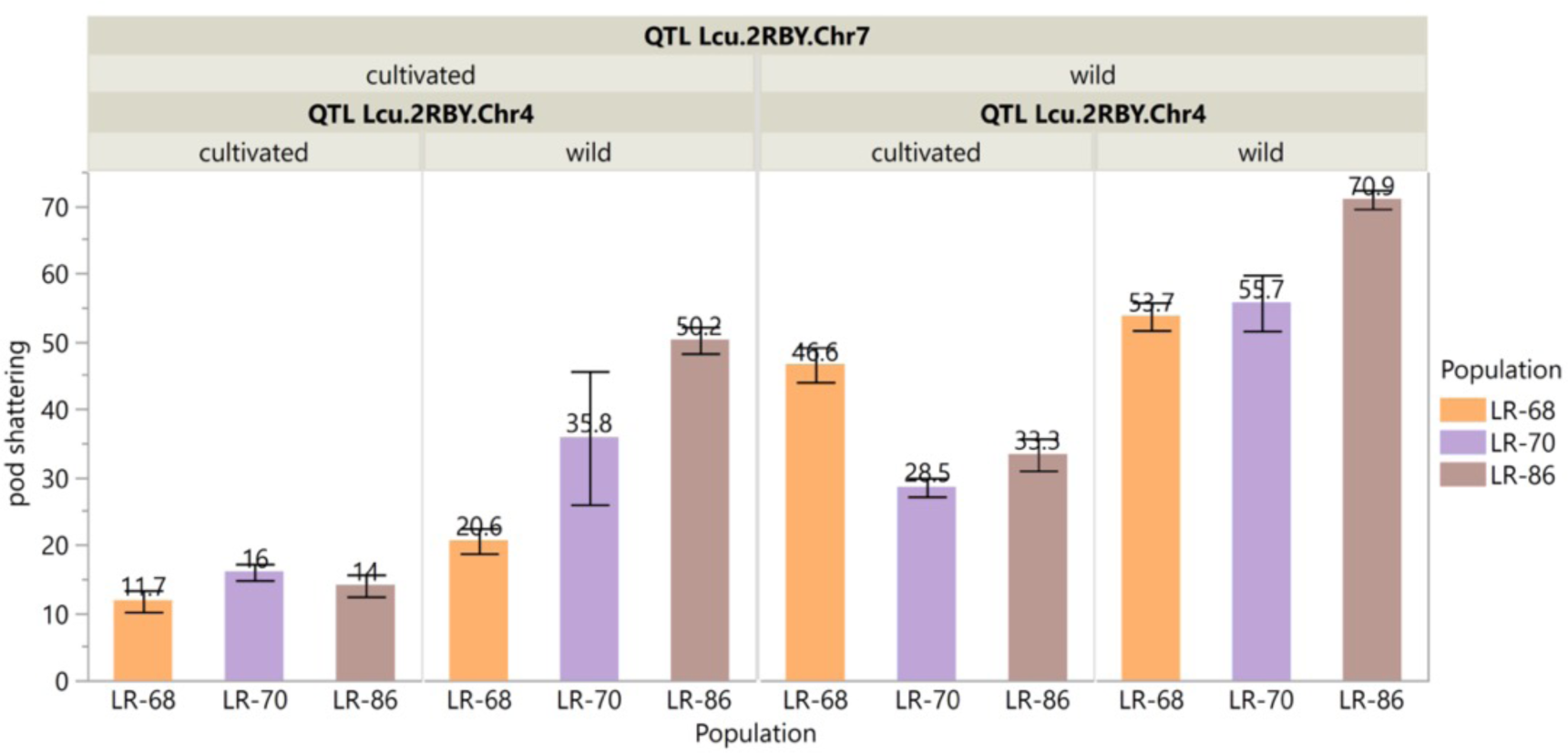
Mean RIL shattering values for genotypes of RILS from LR-68, LR-70, and LR-86 based on their haplotype (wild or cultivated) combination at two consistently detected QTLs on Lcu.2RBY chromosomes 4 and 7. Error bars represent the standard error of the mean.

### Putative genes controlling pod shattering

Our search for potential shattering genes was centered on the regions of Lcu.2RBY chromosomes 4 and 7, where QTLs from LR-68, LR-70, and LR-86 consistently overlapped (Fig. 5; Tables S5). When comparing protein and upstream regulatory DNA sequences of putative genes between cultivated and wild forms, conclusive variations were only observed in the protein coding sequences. For the upstream regulatory DNA sequences, the ultra-high diversity amongst accessions made it challenging to distinguish between cultivated and wild forms with our sample sizes.

One gene, *Lcu.2RBY.4g071110*, within the shared QTL interval on chromosome 4 of Lcu.2RBY, was identified as a shattering gene homolog of *SHP1.A9* (MYB-like transcription factor) in *Brassica napus* (Liu et al. 2006). The corresponding alleles for *Lcu.2RBY.4g071110* in other wild lentil genomes were *Lod.1TUR.4g056470* in Lod.1TUR, *Lor.1WPS.4g061490* in Lor.1WPS, *Ler.1DRO.4g054420* in Ler.1DRO, and *Ler.1DRT.4g076970* in Ler.1DRT. Upon aligning their protein sequences, a variation was detected at positions 54-57 within an aspartic acid-rich domain (Table S6) that was distinct between wild and cultivated accessions. The cultivated protein Lcu.2RBY.4g071110.1 exhibited the sequence “DDGD”, while the wild proteins Lod.1TUR.4g056470.1, Lor.1WPS.4g061490.1, Ler.1DRO.4g054420.1, and Ler.1DRT.4g076970.1 displayed the sequences “D-GD”, “DDAD”, “-DG-”, and “--GD”, respectively.

Within the shared QTL region on chromosome 7 (Fig. 5), three Lcu.2RBY genes are homologs of known shattering genes (Table S5). After comparing the protein sequences between cultivated and wild lentils, we observed consistent sequence variation in *Lcu.2RBY.7g057310* (MADS-box transcription factor). The corresponding protein sequence differed from the protein sequences of the wild alleles at positions 52-57 within a MADS-box domain (Table S6).

## DISCUSSION

Pod shattering negatively impacts recoverable seed yield and can lead to weediness in subsequent seasons, and therefore is typically undesirable in cultivated crops. The extent of pod shattering varies, with modern crop cultivars typically exhibiting a low level, while crop wild relatives display a high degree of shattering. In the process of transferring beneficial alleles (for disease, nutrition, etc.) from wild species to cultivated crops, a primary concern is linkage disequilibrium with shattering alleles, especially if they are closely linked in the genome.

To use wild lentil species more strategically in a breeding program, it is important to understand the genomic location of genes associated with shattering, and the implications of structural variation in the wild species relative to the cultivated species and their effects on pairing and recombination. This information will allow the breeder to design strategies to avoid pod shattering-related alleles during the introgression of desirable wild alleles. In this study, pod shattering was examined in three different interspecific lentil populations to identify QTL regions containing genes responsible for the regulation of seed shattering. Our primary objective was to know whether these shattering-related QTLs could be consistently mapped across different genetic backgrounds and determine the extent of linkage drag in chromosomal segments that are associated with them.

Broad-sense heritability for percent pod shattering varied among the populations, but this may be more related to the growing environments sampled than to the actual population. The higher heritability (*H*^2^ = 77%) observed in LR-68 may be due to the two research sites being grown within the same year, leading to restricted environmental variation. The moderate heritability estimates for LR-70 (*H*^2^ = 46%) and LR-86 (*H*^2^ = 54%) were derived from multiple locations across three years (2016, 2017, and 2018) capturing a higher level of environmental variability. For instance, there was significant variation in rainfall during those growing season (April – September), ranging from 112 mm in 2018 to 244 mm in 2016 (https://saskatoon.weatherstats.ca/charts/precipitation-monthly.html).

Intrachromosomal translocations in one parent relative to the other led to pseudolinkage and reduced recombination in all three populations. In LR-70, reciprocal translocations between chromosomes 1 and 5, and among chromosomes 2, 6, and 7 of the wild *L. odemensis* parent relative to the cultivated *L. culinaris* parent resulted in extensive pseudolinkage and only four linkage groups in the map (Fig. 4). These regions were also all associated with a reduction in recombination leading to large stretches of binned markers and resulting in a shrinkage in the size of the genetic linkage map in this population relative to the other two populations.

A chromosome 2-7 reciprocal translocation in *L. orientalis* relative to *L. culinaris* (Fig. 5) similarly caused pseudolinkage between markers from these chromosomes in the LR-86 genetic linkage map (Figure 5). Interestingly, the pseudolinkage in the LR-68 (*L. orientalis x L. culinaris*) map occurred between markers from chromosomes 2 and 5 (Figure 3) indicating that there are likely multiple patterns of chromosomal rearrangements between *L. orientalis* and *L. culinaris*, and that the current *L. orientalis* genome assembly (Lor.1WPS; the parent of LR-86) is not representative of the whole species. This is supported by an earlier study by Dissanayake et al. (2020), who found that *L. orientalis* is divided into at least two distinct clusters; one being closer to *L. culinaris* and the other clustering more closely with *L. odemensis*, indicating that different *L. orientalis* populations may have undergone separate evolutionary processes.

Even in the linkage groups associated with chromosomes not involved in large, interchromosomal translocations, there were regions of reduced recombination as evidenced by the binning of large numbers of markers. For instance, two bins in LG6 of LR-86 at 30-31 cM containing 83 markers and another two bins in LG6 of LR-68 at 93-94 cM consisting of 40 markers, both covering more than 20 MB. Genes in these regions that are linked in repulsion with the shattering allele will be difficult to separate using common breeding methods and will require, at a minimum, increased population sizes or possibly other strategies to break the linkage.

Across the three populations, 15 significant QTLs were identified but several were clustered consistently on chromosomes 4 (434 to 442 MB) and 7 (437 to 473 MB). Identifying multiple QTLs contrasts with the previous findings from Chen (2018), Ladizinsky (1979), and Tahir and Muehlbauer (1994), who reported that lentil dehiscence was controlled by one or a few genes. This is likely because they used binary scales to score pod shattering, which does not capture the variability in percentage of shattered pods per plant that we noted. Cultivated lentil pods will shatter if exposed to some environmental conditions, contributing in a small way to the observed variability. Our QTL results do, however, show one major locus on chromosome 7 does contribute upwards of 22 (LR-70) to 40 % (LR-86) to the variability in this trait. Chen (2018) identified a major locus in a similar region on chromosome 7 in a RIL population (LR-26) developed from *L. culinaris* cv. ‘Eston’ x *L. ervoides* acc. IG 72815. This QTL on chromosome 7 is likely the DH (dehiscence) locus with large effect on LGIII of the Fratini et al. (2007) genetic map generated using the F_2_ population that led to the LR-86 RILs. Fratini et al. (2007) noted that this locus was not located in the linkage groups associated with cotyledon color (*Yc*), brown seed coat color (*Ggc*), or seed coat pattern (*Scp*), which have been respectively mapped on chromosomes 1, 2, and 6 (Subedi et al. 2019), further suggesting it is the same one. This locus could be also the *Pi* gene, identified as a monogenic gene controlling lentil pod shattering by Tahir and Muehlbauer (1994). Previously, Vaillancourt and Slinkard (1993) stated that the *Pi* locus was not linked to tan seed coat color (*Tgc*), which is on chromosome 3, or to *Yc* on chromosome 1 (Subedi et al. 2019). They also believed it was unlikely to be linked to *Scp*. Then *Pi* should not be on chromosomes 1, 3, or 6, again suggesting it could be the major gene on chromosome 7. Other studies, however, have indicated that *Pi* is in a region syntenic to the pea gene *Dpo1*, implying that these two genes might be homologous (Weeden et al. 1992, 2002). The *Dpo1* is situated on pea chromosome 3, which corresponds to lentil chromosome 6 (Weeden et al. 1992; Hradilová et al. 2017; Ramsay et al. 2021). This creates a contradiction, as we found no major QTL on that chromosome. Another suggestion is that *Pi* is MACE-P015, which aligns with Medtr2g079050 (Hradilová et al. 2017) and is homologous to the lentil gene Lcu.2RBY.2g080190 on lentil chromosome 2. Our QTL mapping only revealed a few minor QTLs on Lcu.2RBY chromosome 2, which are unlikely to be this gene.

Another locus that was consistently detected across the three populations was on Lcu2.RBY chromosome 4. It has a lesser effect and might correspond to the gene with an intermediate effect that Fratini et al. (1987) identified on their LGV. This LG also did not associate with *Yc* (on chromosome 1), *Scp* (on chromosome 6), or *Ggc* (on chromosome 2), leaving chromosome 4 as a strong possibility.

Annotated lentil genes within the consistent QTL intervals on Lcu.2RBY chromosomes 4 and 7 were queried to determine if any of them exhibited homology to known pod shattering genes in other plant species. Following a protein sequence alignment of these matches in the cultivated Lcu.2RBY genome against their corresponding alleles in wild lentil genomes, we identified two potential pod shattering candidates. *Lcu.2RBY.4g071110*, located on the distal segment of Lcu.2RBY chromosome 4, encodes a MYB-like transcription factor and is a homologue of *SHP1.A9* which regulates the development and cell wall degradation in *Brassica napus* (Li et al. 2006). This gene also corresponds to *SHAT1-5* that causes pod shattering in soybean via altering the thickness of lignified fiber cap cells in the abscission layer (Dong et al. 2014). The other candidate gene is *Lcu.2RBY.7g057310* on Lcu.2RBY chromosome 7. It encodes a MADS-box transcription factor that shares homology with the *SH8* gene in rice, which is recognized for controlling seed shattering, although its specific mode of action has yet to be elucidated (Li et al. 2006).

Changes to the coding region of a candidate gene is not the only known variation that has been linked to shattering. Changes in upstream regulatory sequences were demonstrated by Dong et al. (2014), who found a 20 bp deletion in the promoter region that hindered the expression of *SHAT1-5* and caused pod shattering in soybean. We found *Lcu.2RBY.7g056560* a homologue of *SHAT1-5* within the QTL on Lcu.2RBY chromosome 7. Due to a limited sample size and high diversity of upstream sequences, however, our alignment of the promoter region between cultivated and wild sequences remains inconclusive for this gene. Moving forward, a revisit of these candidate genes using time-course gene expression analysis, along with extensive sequencing of the upstream promoter regions in both cultivated and wild lentils, is recommended to further confirm their candidacy.

In conclusion, we have identified two key regions of the wild lentil genomes that we should be prepared to select against when introgressing beneficial loci into cultivated lentil. Having markers could greatly facilitate this effort. Avoiding the locus on chromosome 4 should be reasonably simple as there are no observed translocations, and pairing and recombination of the chromosomes should be normal. The one on chromosome 7, however, will prove a bit more challenging in the interspecific combinations where there is a translocation in one genome relative to the other. These translocations lead to multiple sets of chromosomes pairing during meiosis which typically leads to duplications and deletions and reduced fertility. The gametes that survive typically retain the parental chromosomes of the pairs with few to no recombinations. If the beneficial allele is not on one of these chromosomes this is not an issue, but if it is, then it will be difficult to break up linkage blocks. Having a detailed understanding of the chromosomal make-up of the genomes involved in interspecies hybrids will assist in the design of crossing strategies to mine wild relatives for beneficial alleles.

## Abbreviations

LG: linkage group
QEI: QTL by environment interaction
QTL: quantitative trait locus
RIL: recombinant inbred line
SNP: single nucleotide polymorphism
VCF: variant call format

## ACKNOWLEDGEMENTS

The study was started under the ‘Application of Genomics to Innovation in the Lentil Economy (AGILE)’ project and completed under the ‘Enhancing the Value of Lentil Variation for Ecosystem Services (EVOLVES)’ project, both funded by Genome Canada and managed by Genome Prairie. We express our gratitude for the financial support provided by Saskatchewan Pulse Growers, Western Grains Research Foundation, the Government of Saskatchewan, and the University of Saskatchewan, which matched the funding received. We would like to thank Brent Barlow and his Pulse Crop Breeding team for technical support, and to Robert Stonehouse, Larissa Ramsay, and Hai Ying Yuan for providing genomic data.

## DATA AVAILABILITY

Genetic linkage maps and phenotypic data and metadata are available via: https://knowpulse.usask.ca/study/shattering-loci-case-study. Additional analysed data are available as supplemental data.

## SUPPLEMENTAL MATERIAL

Fig. S1. Distributions of square-root transformed pod shattering results for lentil interspecific RIL populations LR-68, LR-70, and LR-86 across six environments. Black vertical lines indicate values for the wild parent while red vertical lines indicate the cultivated parent in each environment.

Fig. S2. Pearson’s correlation coefficients and broad-sense heritability estimates (*H*^2^) on a multi-environment basis of pod shattering in lentil interspecific populations LR-68 (A – two environments), LR-70 (B – three environments), and LR-86 (C – 5 environments). Estimation of proportion of variation explained by genotype, environment, genotype × environment, and residue (D). The proportion of variation explained by genotype was used to represent *H*^2^.

Fig. S3. Pairwise recombination frequency and LOD scores for all pairs of markers in genetic linkage maps of LR-68 (A), LR-70 (B), and LR-86 (C). Estimates are illustrated by a heatmap with dark blue indicating no linkage (low LOD and high recombination frequency) and yellow signifying tight linkage (high LOD and low recombination frequency).

Table S1. Sequences of 48 known pod shattering proteins from various plant species obtained from NCBI database.

Table S2. Summary of pod shattering (%) of parents and their recombinant inbred line (RIL) progeny in LR-68, LR-70, and LR-86 populations across environments.

Table S3. Pod shattering (%) variance component estimation of genotype, environment, and genotype × environment in LR-68, LR-70, and LR-86 populations.

Table S4. Summary statistics of 15 pod shattering QTLs in LR-68, LR-70, and LR-86 populations.

Table S5. Lcu.2RBY genes with their known shattering gene homologs within QTL regions identified in LR-68, LR-70, and LR-86 populations.

Table S6. Alignment of protein sequences from candidate genes from the cultivated Lcu.2RBY genome with their corresponding alleles in wild lentil genomes.

