## Supplemental Figures for "Understanding Genome Structure Facilitates the Use of Wild Lentil Germplasm for Breeding: A Case Study with Shattering Loci"

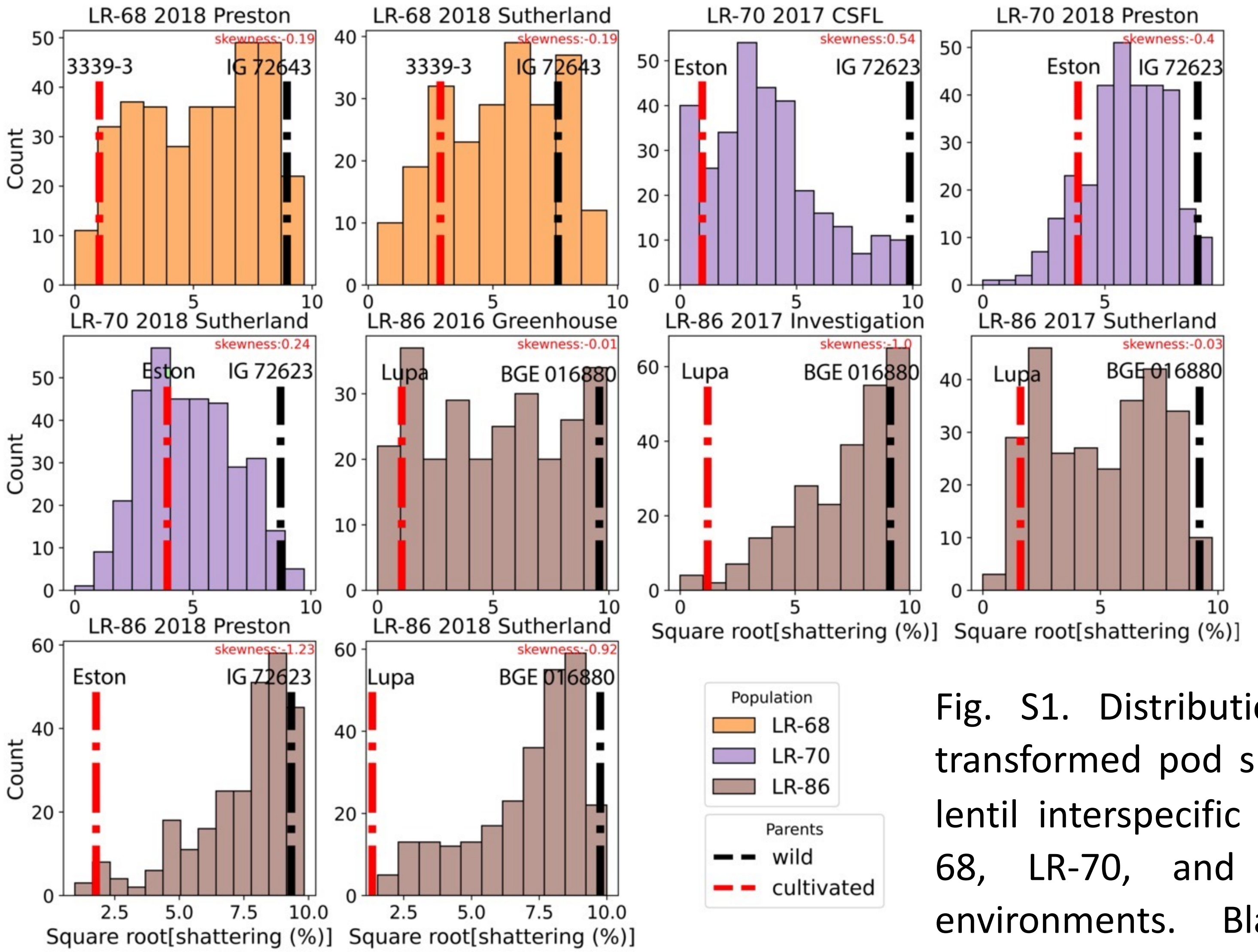


Fig. S1. Distributions of square-root transformed pod shattering results for lentil interspecific RIL populations LR-68, LR-70, and LR-86 across six environments. Black vertical lines indicate values for the wild parent while red vertical lines indicate the cultivated parent in each environment.

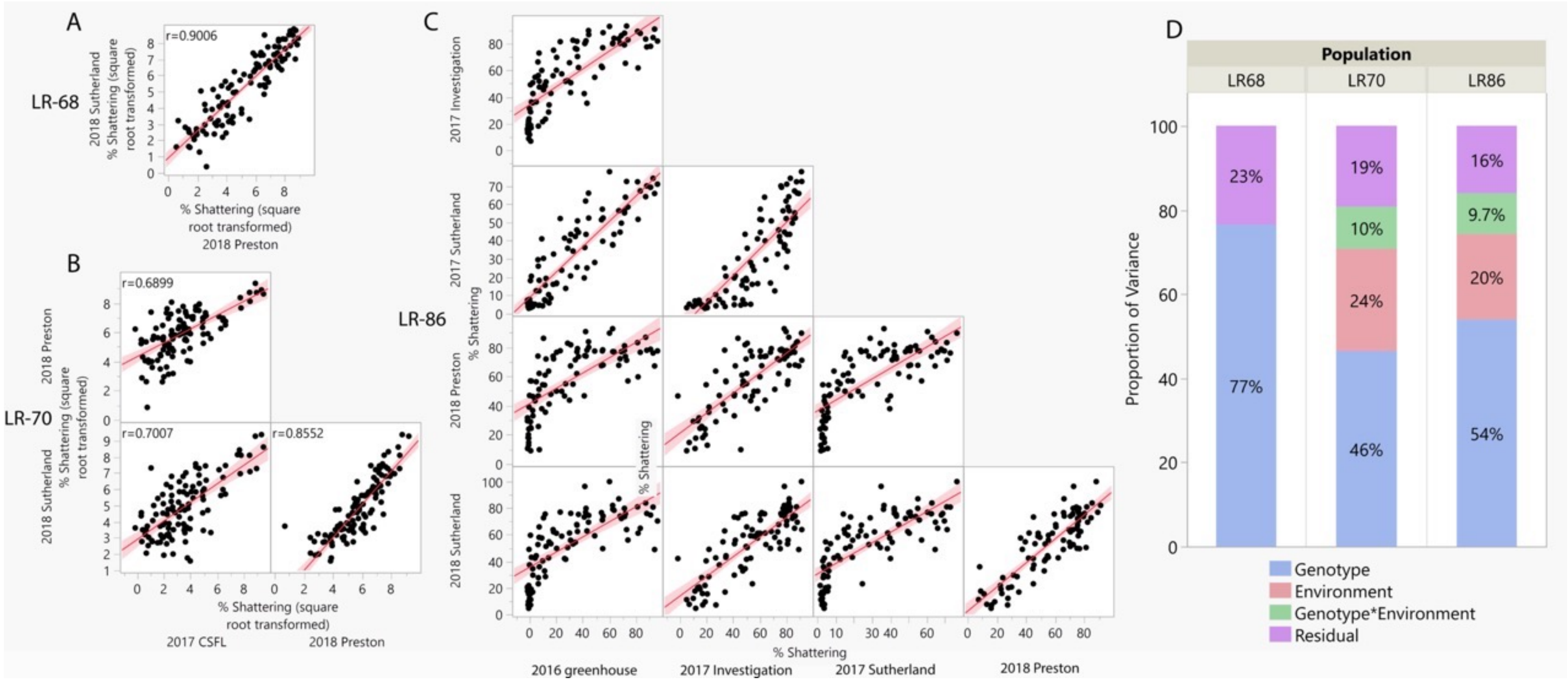


Fig. S2. Pearson's correlation coefficients and broad-sense heritability estimates ($H^2$) on a multi-environment basis of pod shattering in lentil interspecific populations LR-68 (A – two environments), LR-70 (B – three environments), and LR-86 (C – 5 environments). Estimation of proportion of variation explained by genotype, environment, genotype $\times$ environment, and residue (D). The proportion of variation explained by genotype was used to represent $H^2$.

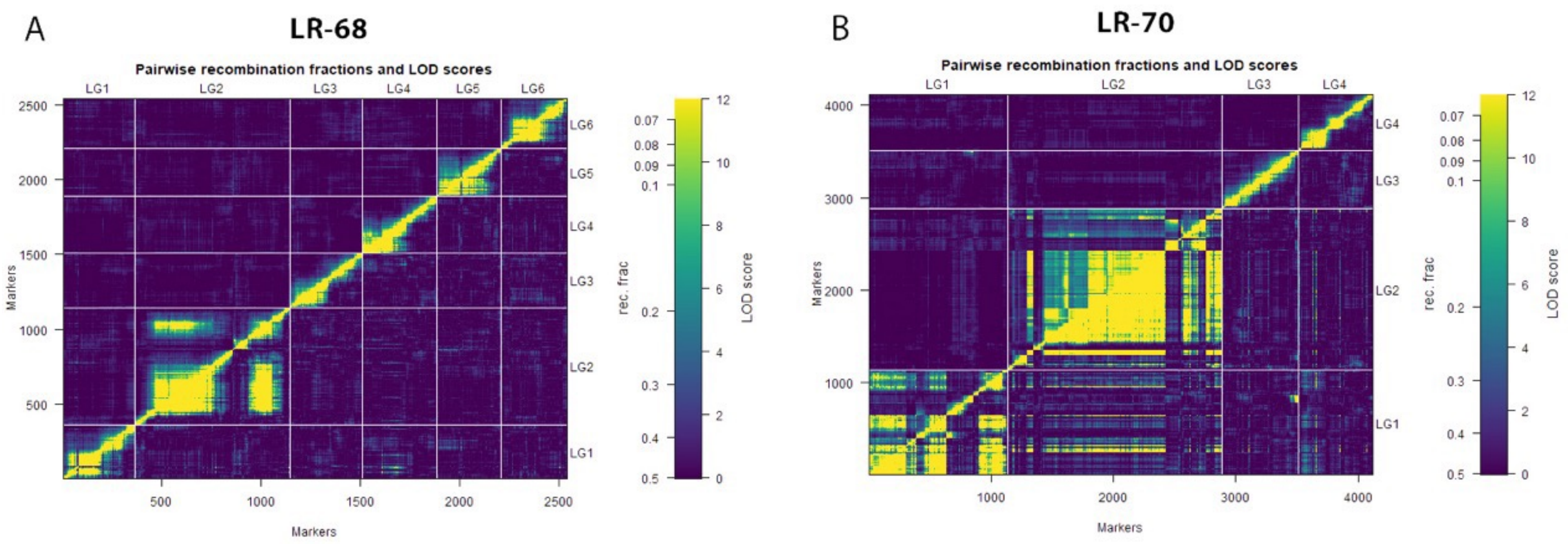


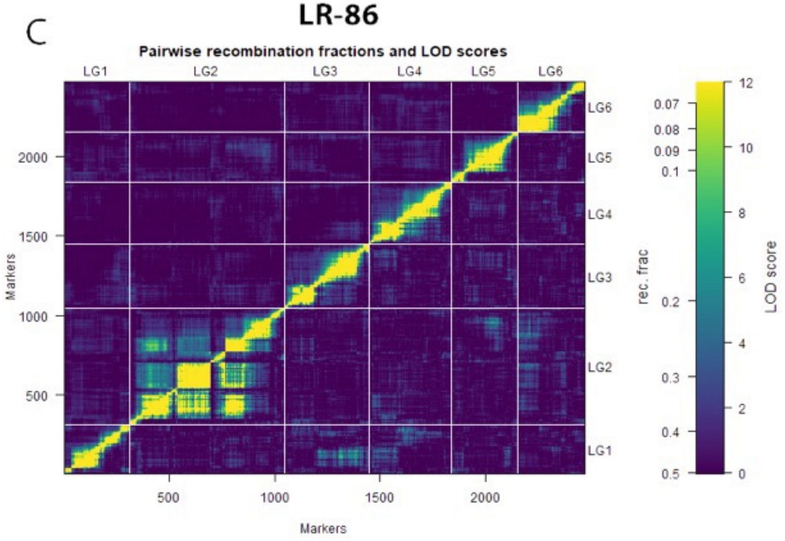


Fig. S3. Pairwise recombination frequency and LOD scores for all pairs of markers in genetic linkage maps of LR-68 (A), LR-70 (B), and LR-86 (C). Estimates are illustrated by a heatmap with dark blue indicating no linkage (low LOD and high recombination frequency) and yellow signifying tight linkage (high LOD and low recombination frequency).
